# Network Analysis Reveals Synergistic Genetic Dependencies for Rational Combination Therapy in Philadelphia Chromosome-like Acute Lymphoblastic Leukemia

**DOI:** 10.1101/2021.01.06.425608

**Authors:** Yang-Yang Ding, Hannah Kim, Kellyn Madden, Joseph P Loftus, Gregory M Chen, David Hottman Allen, Ruitao Zhang, Jason Xu, Yuxuan Hu, Sarah K Tasian, Kai Tan

## Abstract

Systems biology approaches can identify critical targets in complex cancer signaling networks to inform therapy combinations and overcome conventional treatment resistance. Herein, we developed a data-driven, network controllability-based approach to identify synergistic key regulator targets in Philadelphia chromosome-like B-acute lymphoblastic leukemia (Ph-like B-ALL), a high-risk leukemia subtype associated with hyperactive signal transduction and chemoresistance. Integrated analysis of 1,046 childhood B-ALL cases identified 14 dysregulated network nodes in Ph-like ALL involved in aberrant JAK/STAT, Ras/MAPK, and apoptosis pathways and other critical processes. Consistent with network controllability theory, combination small molecule inhibitor therapy targeting a pair of key nodes shifted the transcriptomic state of Ph-like ALL cells to become less like kinase-activated *BCR-ABL1*-rearranged (Ph+) B-ALL and more similar to prognostically-favorable childhood B-ALL subtypes. Functional validation experiments further demonstrated enhanced anti-leukemia efficacy of combining the BCL-2 inhibitor venetoclax with tyrosine kinase inhibitors ruxolitinib or dasatinib *in vitro* in human Ph-like ALL cell lines and *in vivo* in multiple patient-derived xenograft models. Our study represents a broadly-applicable conceptual framework for combinatorial drug discovery, based on systematic interrogation of synergistic vulnerability pathways with pharmacologic targeted validation in sophisticated preclinical human leukemia models.

## INTRODUCTION

Cancer cells exploit multiple deregulated pathways to evade the selective pressure of single-agent drugs, promoting therapeutic resistance and clinical relapse^1^. However, combination therapy regimens for cancer have traditionally been non-specific with broad toxicity profiles and developed in an *ad hoc* manner. More rational identification of new targets in human cancers for combination drug regimens is an essential next step. There is growing interest in identifying synergistic genetic interactions as targets for combination therapy^2^, but large-scale experimental screening for genetic interactions has been technically challenging and expensive given the large number of candidate gene pairs one has to screen. As a result, existing RNA-interference and CRISPR-based screenings have been limited to only a few hundred genes^3,4^, far from saturating the search space of all possible (~4×10^8^) pairwise interactions in the human genome. Given the above challenges, we developed a systems biology approach that enables efficient *in silico* genetic screening and prioritization of co-targetable pathways for combinatorial therapeutics followed by rigorous *in vitro* and *in vivo* pharmacologic validation in a difficult-to-cure subtype of leukemia.

Philadelphia chromosome-like acute lymphoblastic leukemia (Ph-like ALL) comprises 15-30% of high risk B-ALL cases in children and adolescents/young adults (AYAs) and 20-40% in older adults^5–9^, and is associated with high rates of conventional chemotherapy resistance and poor clinical outcomes^9,10^. Ph-like ALL is defined by a kinase-activated transcriptomic signature resembling that of Philadelphia chromosome-positive (Ph+) ALL, but lacks the *BCR-ABL1* rearrangement^11^. Ph-like ALL is instead driven by alternative genetic alterations in two major subclasses: (1) JAK/STAT pathway alterations involving *CRLF2, JAK2, EPOR, IL7R,* or *SH2B3* rearrangements or indels and (2) ABL-class kinase fusions involving *ABL1, ABL2, CSF1R,* or *PDGFRB* rearrangements^10^. Preclinical studies of tyrosine kinase inhibitor (TKI) monotherapy in Ph-like ALL models have expectedly demonstrated incomplete anti-leukemia activity^12–15^ likely via compensatory signaling mechanisms, emphasizing the need for more rationally-designed combination therapy approaches to achieve cure. In the present studies, we hypothesized that an unbiased systems biology approach could effectively elucidate optimal target pairings. Our network-based analysis is optimal to address the unique challenges of Ph-like ALL given its known dysregulation of multiple intracellular pathways that maintain a high degree of crosstalk.

A main goal of effective multi-agent therapy is identifying drug combinations with synergistic efficacies, but not synergistic toxicities. We recently reported our OptiCon (<u>Opti</u>mal <u>Con</u>trol) algorithm^16^ that is capable of discovering novel disease-specific synergistic regulators by integrating a molecular interaction network with large-scale patient genomic and transcriptomic data. OptiCon is based on the theory of network controllability, a mathematically validated framework for identifying a set of driver nodes in a complex network that can guide the system from an initial state to any desired final state^17^. OptiCon integrates clinical, genomic and expression data, as well as a gene regulatory network to identify critical network nodes termed Optimal Control Nodes (OCNs) that control a maximal number of deregulated genes (for optimal therapeutic efficacy) and a minimal number of unperturbed genes (for toxicity minimization). Synergistic regulators (OCN pairs) are then nominated based on the synergy score, which quantifies the degree of crosstalk between pathways downstream of the two OCNs and the amount of enrichment for deregulated and mutated genes in the Optimal Control Regions (OCRs) of the two OCNs^16^. A high proportion of the gene pairs nominated by this algorithm has known synthetic lethal genetic interactions^16^. In the current study, we leveraged this powerful computational tool to identify key oncogenic dependencies in Ph-like ALL and to prioritize pathways for pharmacologic targeting *in vitro* and *in vivo* using human cell lines and various preclinical patient-derived xenograft (PDX) models of *CRLF2*-R or ABL-class Ph-like ALL.

## METHODS

Detailed experimental methodologies are included in the Supplemental Data.

### Prediction of candidate combination therapeutic targets

We used our OptiCon^16^ algorithm to computationally nominate candidate combination therapeutic targets using the following inputs: a gene regulatory network, leukemia-specific genetic mutation data, and gene expression data (Tables S1-S4). Whole genome sequencing (WGS), whole exome sequencing (WES), and microarray expression data from B-ALL samples generated by the Therapeutically Applicable Research to Generate Effective Treatments (TARGET) project^18^ and the Pediatric Cancer Genome Project (PCGP)^7^ were used for mutation calling and differential expression analyses.

### Cell lines

Ph-like ALL cell lines MUTZ5 and MHH-cALL4 were obtained from the DSMZ cell biorepository. Ph-like TVA-1 cells with *ETV6-ABL1* fusion were immortalized *in vitro* as a cell line from a PDX model established by the laboratory of Dr. David Fruman at the University of California, Irvine^19^.

### Drug synergy testing *in vitro*

Cell lines were incubated at drug concentrations ranging between 1 nM - 50 μM for 72 hours. Cell viability was assessed by Cell-Titer Glo viability assays (Promega). IC_50_ values were determined using GraphPad Prism. Combination index (CI) values were calculated using Compusyn software^20^.

### RNA sequencing and transcriptome analysis

Cells from the three Ph-like cell lines were cultured in medium containing vehicle, TKI alone, venetoclax alone, or in combination at indicated concentrations for 72 hours. RNA was extracted and sequenced using established protocols. See supplemental methods for full details of sequencing protocols and transcriptome analyses by differential gene expression, dimensionality reduction, and gene signature score methods.

### Immunoblotting

Protein and phospho-protein levels were measured by Western blot analysis using standard protocols. Antibodies used are described in supplemental methods.

### Patient-derived xenograft modeling and *in vivo* drug combination trials

Patient-derived xenografts of primary patient leukemia samples were established as previously described^12,13^. Viably cryopreserved diagnostic bone marrow ALL cells from patients were engrafted into NOD.Cg-Prkdcscid Il2rgtm1Wjl/SzJ (NSG) mice via informed consent on Institutional Review Board- and Institutional Animal Care and Use Committee (IACUC)-approved research protocols. See supplemental methods for further details.

### Data Sharing Statement

RNA sequencing data are available at GEO. All software supporting the analysis in this study can be found in public repositories. Software package implementing OptiCon has been deposited at GitHub (https://github.com/tanlabcode/OptiCon).

## RESULTS

### Network-controllability analysis of patient multi-omics datasets identifies targetable synergistic regulators in the Ph-like ALL gene network

We applied our unbiased OptiCon algorithm to the study of Ph-like ALL with an overarching goal of identifying synergistic target pairings for biologically-rational combination therapy (Figure 1A). We analyzed WGS, WES, and gene expression microarray data from 1046 primary childhood/AYA B-ALL specimens, of which 289 were Ph-like (Table S1). We identified structural variants, small indels, and point mutations (Tables S2-S3) and differentially expressed genes in Ph-like B-ALL as compared to prognostically-favorable B-ALL subtypes that are highly curable with conventional chemotherapy, including the *ETV6-RUNX1*^21^, high hyperdiploidy^22^, and *DUX4*-rearranged/*ERG*-dysregulated subtypes^23^, in order to elucidate the key genetic dependencies that may render Ph-like ALL less sensitive to conventional chemotherapy. Using these data and a high-quality curated gene regulatory network (Table S4) as inputs, OptiCon analysis predicted 81 key regulator gene pairs to be significantly synergistic in Ph-like ALL (synergy score p-values <0.05, Table S5), which represent specific pairings of 14 OCNs (Table 1, Figure S1-B).

**Figure 1.**
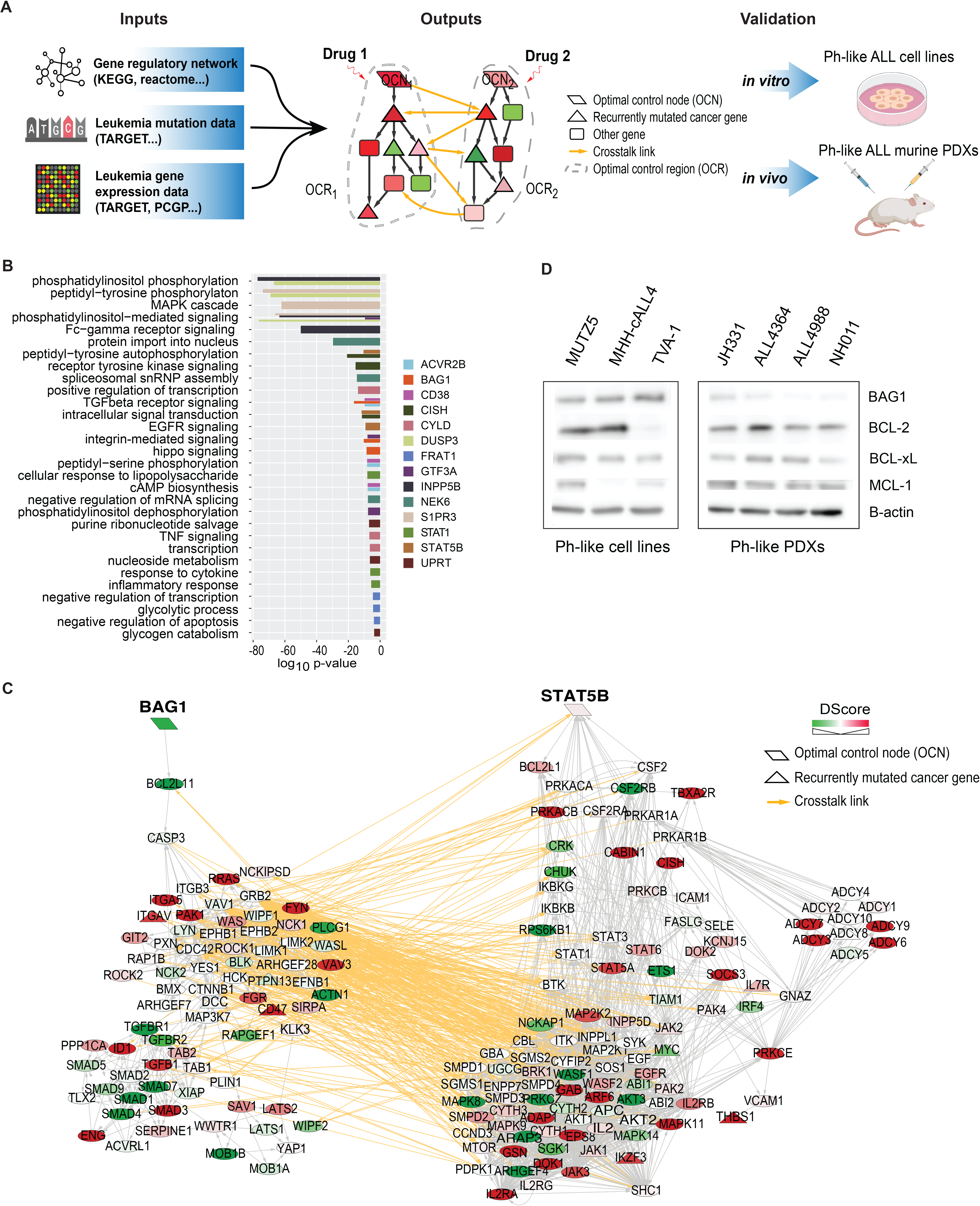
A systems biology approach to discovery and testing of combinatorial therapeutic targets. **(A)** Overview of the Optimal Control (OptiCon) network-based approach towards identifying and validating synergistic drug targets in Ph-like B-ALL. OptiCon input data include a high-quality curated gene regulatory network, genetic mutation data, and gene expression data under two conditions (Ph-like ALL versus favorable-risk B-ALL subtypes). Output of OptiCon identifies synergistic Optimal Control Node (OCN) pairs. Druggable pathways defined by OCNs and their respective Optimal Control Regions (OCRs) are then validated *in vitro* in Ph-like ALL cell lines and *in vivo* in murine patient-derived xenograft (PDX) models. **(B)** Enriched Gene Ontology (GO) terms among predicted OCRs. Each color represents the OCR of a predicted OCN. Thickness of bars varies since certain terms were enriched among multiple OCRs. **(C)** Synergistic OCN pair S*TAT5B* and *BAG1* predicted for Ph-like ALL. Gene crosstalk links between their specific OCRs are shown in yellow. Shade of a node represents the deregulation score (DScore) of the corresponding gene. Red, up-regulated in Ph-like ALL; green, down-regulated in Ph-like ALL**. (D)** Baseline protein expression of BAG1 and the anti-apoptotic BCL2-family proteins BCL2, BCL-xL, and MCL1 in several Ph-like cell lines and PDXs.

**Table 1.** Optimal control nodes (OCNs) predicted to be synergistic key regulators in Ph-like ALL and their known small molecule inhibitors.

| OCN | Name | Drug/compound inhibitor | Source |
| --- | --- | --- | --- |
| *ACVR2B* | activin A receptor type 2B | bimagrumab | DGIdb |
| *BAG1* | BCL2 associated athanogene 1 | (2R,3R,4S,5R)-2-[6-amino-8-[(3,4-dichlorophenyl)methylamino]purin-9-yl]-5-(hydroxymethyl)oxolane-3,4-diol | DrugBank |
| *CD38* | CD38 molecule | daratumumab, isatuximab, MOR-202, SAR-650984 | DGIdb, TTD, DrugBank |
| *CISH* | cytokine inducible SH2 containing protein | epoetin alfa | DGIdb |
| *CYLD* | CYLD lysine 63 deubiquitinase | NA | NA |
| *DUSP3* | dual specificity phosphatase 3 | NA | NA |
| *FRAT1* | FRAT1, WNT signaling pathway regulator | NA | NA |
| *GTF3A* | general transcription factor IIIA | NA | NA |
| *INPP5B* | inositol polyphosphate-5-phosphatase B | D-Myo-Inositol-1,4-Bisphosphate | DrugBank |
| *NEK6* | NIMA-related kinase 6 | 8205, (5Z)-2-hydroxy-4-methyl-6-oxo-5-[(5-phenylfuran-2-yl)methylidene]-5,6-dihydropyridine-3-carbonitrile | DGIdb, PubChem |
| *S1PR3* | sphingosine-1-phosphate receptor 3 | AFD(R), AUY954, EDD7H9, FTY720-phosphate, VPC03090-P, VPC12249, VPC23019, VPC44116, compound 26 | DGIdb, TTD |
| *STAT1* | signal transducer and activator of transcription 1 | AVT-02 UE | DGIdb, TTD |
| *STAT5B* | signal transducer and activator of transcription 5B | Dasatinib, Ruxolitinib | DGIdb, DrugBank |
| *UPRT* | uracil phosphoribosyltransferase homolog | NA | NA |

Pathway enrichment analysis of these Ph-like ALL-specific OCNs and their respective OCR genes showed enrichment in multiple kinase signaling pathways, in regulation of transcription and apoptosis, and in metabolic processes such as glycolysis and nucleoside metabolism (Figure 1B). Importantly, OptiCon identified *STAT5B* and *CISH* (cytokine-inducible SH2-containing protein, a known negative regulator of *STAT5B*) as OCNs, supporting the robustness of our methodology since *STAT5B* is a major known effector in Ph-like ALL^7,24^. Relatedly, phosphatidylinositol 3-kinase (PI3K) and SRC family kinases (*LYN, LCK, SRC)* were also identified in the downstream OCRs of several predicted OCNs in Ph-like ALL (Figure 1C; Table S6). These findings collectively serve as unbiased support for our and others’ prior pharmacologic studies that demonstrated effective targeting of JAK/STAT, PI3K, and SRC kinase pathway signaling in primary Ph-like ALL cells and preclinical models^15,19,25^. Importantly, OptiCon identified novel OCNs not previously known to be targets in Ph-like ALL, including *BAG1* (BCL2-associated athanogene 1), *DUSP3* (dual specificity phosphatase 3), *CD38,* and *NEK6* (never in mitosis gene A-related kinase 6); these genes have all been implicated in other leukemias or in tumorigenesis^26–29^. We further identified several anti- and pro-apoptotic genes, including *BCL2*, *BCL2L11, BAD, BAX,* and *BCL2L1*, in the OCRs of several OCNs in Ph-like ALL (Table S6).

Next, we queried the drug databases TTD^30^, DrugBank^31^, DGIdb^32^, and NIH PubChem to identify known drugs or new chemical compounds that could be used for potential pharmacologic targeting of predicted nodes. Nine of our identified 14 OCNs matched with drug compounds (Table 1). Some agents have only preclinical testing data available in human cancer, such as the recently-described NEK6 inhibitor^33^. Others are more advanced, including the anti-CD38 monoclonal antibody daratumumab^28^ that is FDA-approved for adults with multiple myeloma and under current phase 1/2 clinical study in children with relapsed/refractory leukemias. Amongst the 81 OCN gene pairs predicted to be significantly synergistic, 32 (40%) were found to have known small molecule inhibitors for both members of the pair (hypergeometric test *p*-value < 0.0001, Table S5). Additionally, 559 of the 973 predicted OCR genes (57%) were found to be targets of known drugs or experimental compounds (hypergeometric test *p*-value < 0.0001). Taken together, these results suggest that computationally predicted synergistic regulators and their target pathways in a disease-specific network are valuable sources for identifying novel drug targets.

Since STAT5 signaling is known to be hyperactivated via both JAK and ABL class kinases in Ph-like ALL^7,15^, OptiCon’s nomination of *STAT5B* provided an attractive target for subsequent validation efforts. Several clinical trials are currently investigating the addition of the Janus kinase inhibitor (JAKi) ruxolitinib to treatment of Ph-like ALL patients with *CRLF2* rearrangements and other JAK/STAT pathway alterations (NCT02723994)^34^ or the addition of the SRC/ABL kinase inhibitor (ABLi) dasatinib to therapy for patients with ABL-class alterations (NCT02883049). Given these ongoing trials incorporating single-agent TKIs, we focused on other key regulons predicted to be synergistic with *STAT5B* that may further optimize combination therapy. One OCN pair that ranked as highly synergistic was *STAT5B* and *BAG1* (synergy score 0.023, adjusted *p* value= 0.016). The latter encodes the BCL2-associated athanogene 1 protein that binds to and enhances the anti-apoptotic effect of BCL-2, likely by preventing its degradation^35,36^, although there is not a well-characterized pharmacologic BAG1 inhibitor. This OptiCon pairing and the several other apoptosis-related genes enriched in the OCRs highlight the potential importance of apoptosis pathways in Ph-like ALL and led us to investigate in the studies detailed below the therapeutic potential of venetoclax, a potent and highly-selective BCL-2 inhibitor approved by the US Food and Drug Administration for treatment of adults with relapsed/refractory chronic lymphocytic leukemia^37^ or acute myeloid leukemia (AML)^38^.

### Pharmacologic co-targeting of STAT5 and BCL-2 has synergistic anti-leukemia efficacy in genetically heterogeneous Ph-like ALL cell lines

We next validated our computational predictions *in vitro* in three known Ph-like ALL cell lines that represent both *CRLF2*-R and *ABL*-class alterations, including MUTZ5 (*IGH-CRLF2*, *JAK2* R683G), MHH-cALL-4 (*IGH-CRLF2*, *JAK2* I682F), and TVA-1 (*ETV6-ABL1)*^19^. Principal component analysis (PCA) of gene expression data from these cell lines and primary pediatric B-ALL patient specimens included in the TARGET and PCGP datasets showed that the Ph-like ALL cell lines clustered together with Ph-like and Ph+ ALL patient samples (Figure S1-A) and separate from the other B-ALL subtypes, recapitulating expression signatures of primary Ph-like ALL samples used in the OptiCon analysis. We then investigated the baseline protein expression of BAG1 and anti-apoptotic BCL-2 family members in Ph-like ALL cell lines and PDX models and found that BAG1, BCL-2, BCL-xL, and/or MCL-1 are detected in all tested Ph-like ALL cell lines and PDX cells (Figure 1D).

To evaluate potential synergy *in vitro*, we treated Ph-like ALL cell lines with venetoclax and ruxolitinib (MUTZ5, MHH-cALL-4) or dasatinib (TVA-1) and assessed cell viability. Figures 2A-C show individual drug response curves for each cell line with corresponding half-maximal inhibitory concentration (IC_50_) values. We confirmed high ruxolitinib IC_50_ values for both MUTZ5 and MHH-cALL-4^39^ and low dasatinib IC_50_ for TVA-1. However, limited cell killing was observed at up to 72 hours of drug incubation despite tested TKI doses as high as 30-50 μM (Figure 2A-C). In contrast, all three Ph-like cell lines showed sensitivity to venetoclax with IC_50_ values <800 nM and near-complete cell killing at 72 hours at higher drug doses, suggesting a strong dependency on BCL-2.

**Figure 2.**
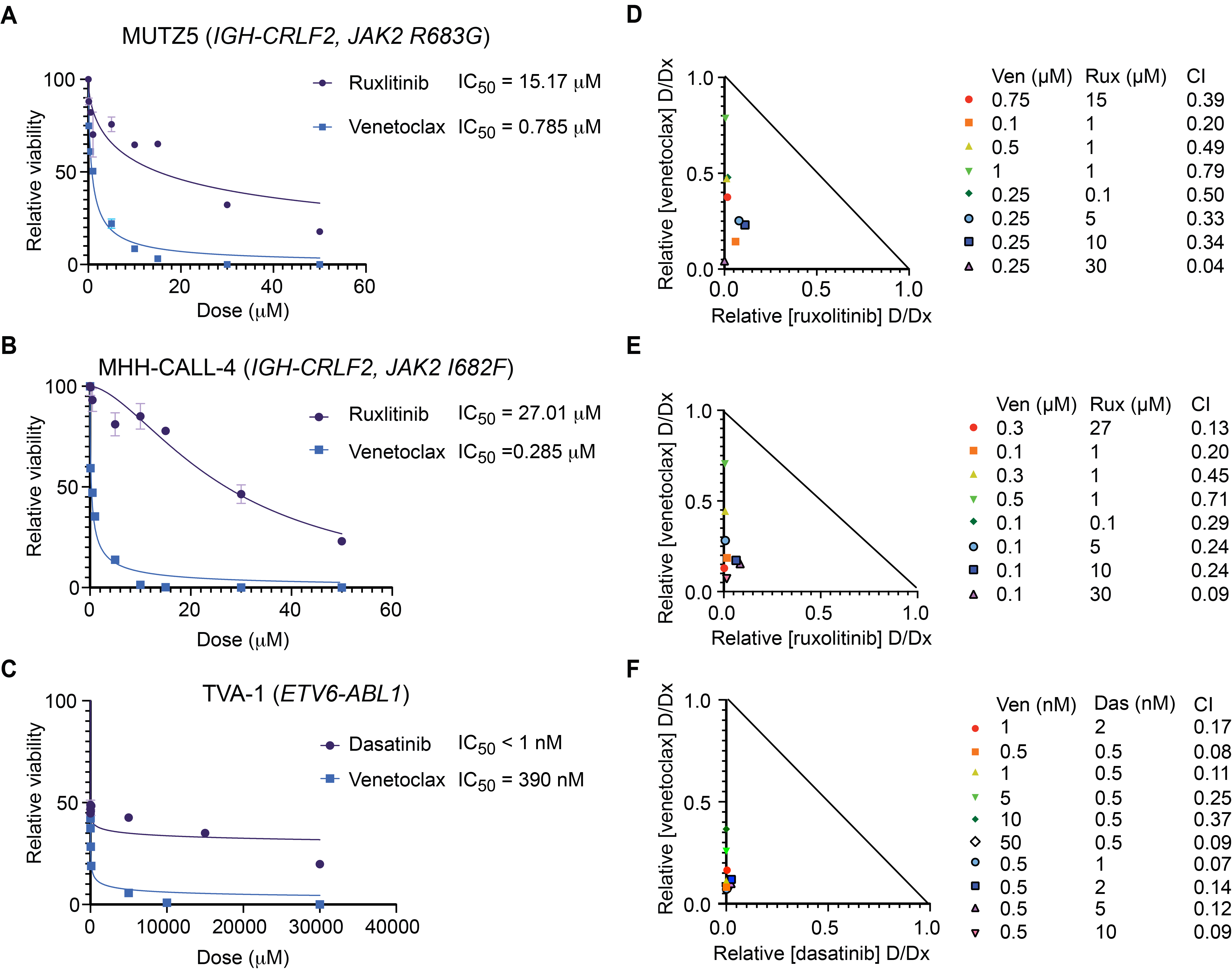
Combined BCL-2 and kinase inhibition is synergistic *in vitro* in Ph-like ALL. Individual dose response curves and IC_50_ values for ruxolitinib (rux) and venetoclax (ven) treatment of **(A)** MUTZ5 and **(B)** MHH-cALL4 cell lines and dasatinib (das) and venetoclax treatment of **(C)** TVA-1 cells. Viability data are shown relative to 0.1% DMSO vehicle assayed at 72 hours using CellTiter-Glo absorbance assays. Each data point represents the mean of six replicate measures +/− standard deviation (SD). **(D-F)** Isobolograms for combination of TKIs with venetoclax at various dose combinations in MUTZ5, MHH-cALL-4 and TVA-1 cell lines, respectively, along with Combination Index (CI) values for each dose combination which were generated using Compusyn. CI values were less than 1.0 (synergistic effect) for all dose combinations tested.

Based on these initial monotherapy IC_50_ data, we then chose a range of drug doses to test in combination and assessed potential drug synergy. Figures 2D-F show the isobolograms and combination index (CI) values of the indicated venetoclax and ruxolitinib or dasatinib combinations; all dosage combinations tested were determined to be significantly synergistic using the Chau-Talalay method (CI<1.0)^20^. Cell viability data with *in vitro* drug combination exposure are shown in Figure S2-A. For subsequent investigations into mechanisms of drug synergy detailed below, we used specific dose combinations as specified in Figure S2-B, chosen because they displayed high synergy (low CI values) in these cell lines (Figure 2D-F), were not completely lethal to the cells, and were not expected to induce appreciable off-target kinase inhibition^40^.

### Combination therapy shifts the transcriptome of Ph-like cells away from Ph+/Ph-like subtypes and towards more chemosensitive B-ALL subtypes

To elucidate the transcriptomic effects of targeted therapy on Ph-like ALL cells, we performed RNA sequencing of the three Ph-like cell lines treated with venetoclax, TKI, or both drugs at our optimized dosing. Analysis of differentially expressed genes (DEGs) and PCA revealed that venetoclax monotherapy effect on the Ph-like ALL transcriptome was minimal (DEGs= 20 using FDR< 0.1, absolute log2FC> 1; Figure S3-A). In contrast, TKI monotherapy and combined TKI and venetoclax exposure had a large, but similar, effect on the transcriptome compared to DMSO control treatment (DEGs= 9651 and 10378, respectively).

We next compared transcriptomes of inhibitor-treated Ph-like ALL cells to those of different B-ALL patient samples using Uniform Manifold Approximation and Projection (UMAP)^41^ to project gene expression data to lower dimensions (Figure 3A). We observed that the transcriptomes of vehicle control-treated and venetoclax monotherapy-treated Ph-like ALL samples, as well as some TKI monotherapy-treated ones, remained similar to those of untreated Ph+ and Ph-like ALL patient samples. Conversely, we found that combined venetoclax and TKI treatment altered the transcriptomic state of Ph-like ALL cells, causing them to cluster closer to B-ALL leukemia subtypes with more favorable cytogenetic alterations. This shift effectively made cells ‘less Ph-like’ and more similar to subtypes that are sensitive to conventional chemotherapy, thereby supporting our network controllability theory (Figure 3B). To quantify further this transcriptome shift in treated Ph-like ALL cells, we developed comparator gene signature scores (Supplemental Methods) for several common B-ALL genetic subtypes. We observed that treating Ph-like ALL cell lines with TKI monotherapy or simultaneous TKI and venetoclax altered their transcriptomes to have higher *ETV6-RUNX1*, hyperdiploidy, and *TCF3-PBX1* gene signature scores (representing favorable risk subtypes) and lower *KMT2A-*rearranged scores (a prognostically-unfavorable subtype^42^) in comparison to control-treated cells (Figure 3C). Therefore, both UMAP and gene signature score analyses support the specific shift of transcriptomic state in Ph-like ALL cells post-treatment.

**Figure 3.**
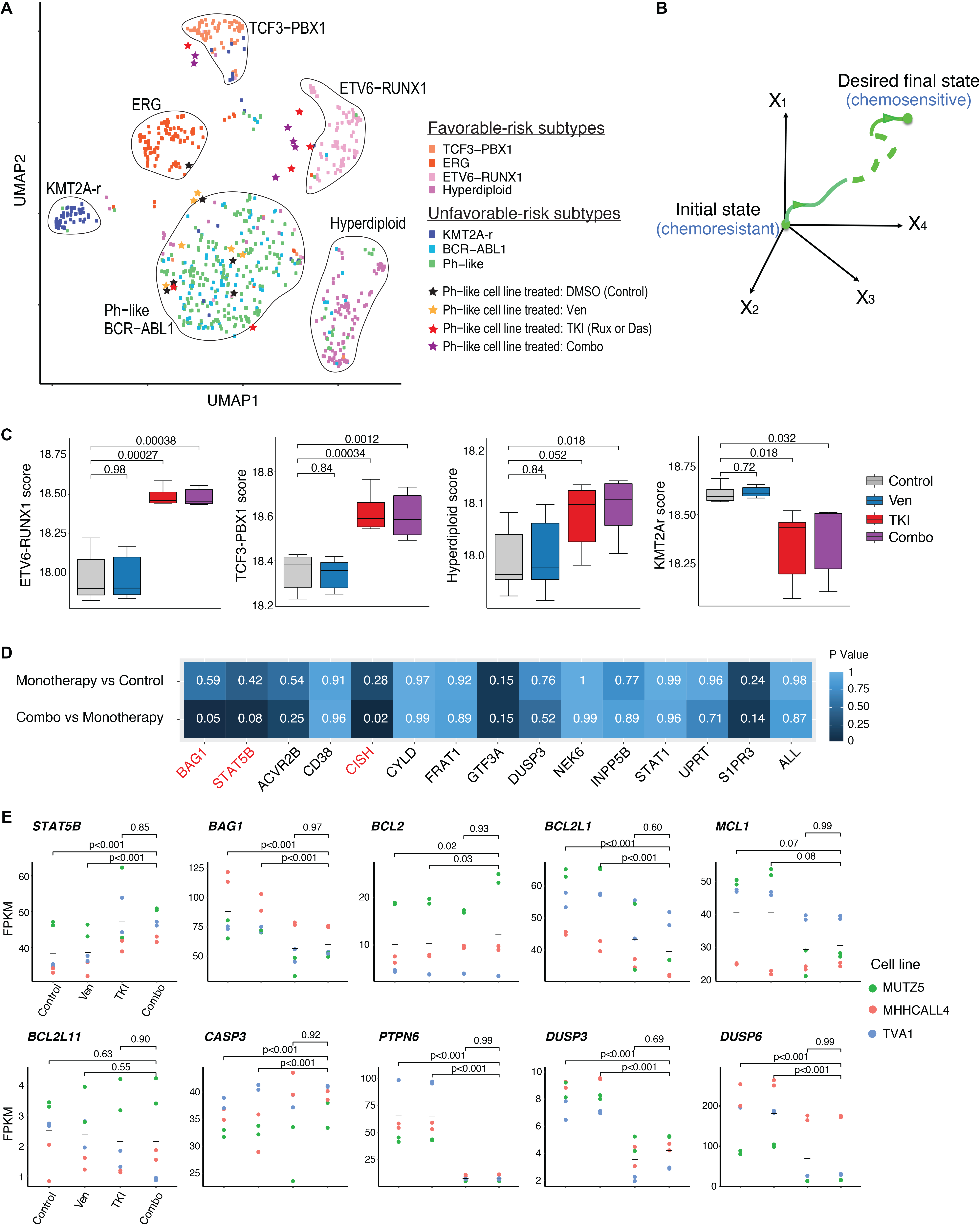
Combination treatment elicits specific and unique changes in the transcriptome of Ph-like cell lines. **(A)** UMAP based on RNA-Seq data from treated Ph-like cell lines and B-ALL patient microarray expression data shows that combination treatment shifts transcriptome of Ph-like cells to resemble that of favorable-risk B-ALL subtypes. **(B)** Network controllability theory posits that one can use control nodes in a gene network to guide a system from an initial state (in this case relatively chemoresistant Ph-like ALL) to a final state (chemosensitive favorable risk subtype B-ALL). **(C)** Signature genes of non-Ph-like B-ALL subtypes (*ETV6-RUNX1*, *TCF3-PBX1,* and hyperdiploidy subtypes) are enriched in Ph-like ALL cells treated with venetoclax and ruxolitinib or dasatinib. Enrichment scores were computed using single-sample Gene Set Enrichment Analysis (ssGSEA). **(D)** Predicted synergistic OCRs *STAT5B* and *BAG1* were significantly perturbed only by combination drug treatment, but not by monotherapy. The degree of perturbation was measured by the overlap between genes in the OCR of each specified OCN and differentially expressed genes in the specified comparison (monotherapy versus control or combination therapy versus monotherapy). Significance of overlap was determined using a hypergeometric test. Significant p-values were observed for the *STAT5B*, *BAG1,* and *CISH* OCRs in dual inhibitor-treated cells, but not with drug monotherapies, and also not in the other OCRs. **(E)** Gene expression changes during monotherapy or combination inhibitor therapy in OptiCon-nominated OCNs and some OCR genes. P-values were calculated using one-way ANOVA implemented in LIMMA software and adjusted for multiple testing using the Benjamini-Hochberg method.

The network controllability theory underlying OptiCon suggests that co-targeting specific OCNs will lead to perturbation of their corresponding OCRs. We thus hypothesized that expression of genes within the identified OCRs of *STAT5B* and *BAG1* would change after combination treatment, but not after monotherapy. Indeed, we observed that *STAT5B* and *BAG1* OCRs were significantly enriched for DEGs only with dual BCL-2 and STAT5 inhibition, but not with single-drug treatment (Figure 3D). On the contrary, none of the OCRs were enriched for DEGs when comparing monotherapy versus control. Interestingly, the OCR of the OCN *CISH* (cytokine inducible SH2 containing protein, a known negative regulator of JAK/STAT signaling) was also enriched for DEGs when comparing dual-inhibitor versus single-agent treatment, although the other 11 OCN-associated OCRs were not appreciably perturbed. Taken together, these findings suggest that Ph-like ALL cells may be shifted towards a potentially more favorable transcriptomic state by direct pharmacologic perturbation of *STAT5B* and *BAG1* and their downstream control regions (Figure 3B).

### Apoptotic and cytokine signaling pathways can be altered by inhibitor therapy, resulting in enhanced apoptosis when combined

We next interrogated the transcriptional, translational, and functional effects of inhibitor therapy on specific pathways nominated by OptiCon. Interestingly, both *BCL2* and *STAT5B* gene expression were upregulated in TKI monotherapy and combined venetoclax/TKI conditions (Figure 3E), which we hypothesize could be due to negative feedback mechanisms such as downregulation of *PTPN6* following TKI treatment^43^. Conversely, other anti-apoptotic genes *BAG1*, *MCL1*, and *BCL2L1* (encoding BCL-xL) were significantly downregulated in combination drug-treated cells versus vehicle control. We also detected significant expression changes in several other genes involved in the PI3K/Akt/mTOR, Ras/mitogen-activated protein kinase (MAPK) pathways, and intrinsic and effector apoptosis mechanisms following *in vitro* inhibitor exposure (Figure S3-B).

On a post-translational level, ruxolitinib or dasatinib treatment was sufficient to completely abrogate activated pSTAT5 and MAPK targets pERK and/or pJNK (Figure 4A-B). Unexpectedly, BCL-2 expression was observed to be highest in venetoclax-treated and combination drug-treated conditions, which could potentially be interpreted as a survival advantage of BCL-2 overexpressing leukemia cells and elimination of low-BCL-2 cells, as has been reported in AML 44, although other studies have shown that elevated BCL-2 expression is not a reliable biomarker of venetoclax sensitivity or resistance^44,45^. MCL-1 on the other hand, is an anti-apoptotic BCL-2 family protein that is tightly transcriptionally regulated and whose high expression is known to mediate venetoclax resistance by binding to BIM^46^. Thus, we noted with interest that levels of the anti-apoptotic proteins MCL-1 and BAG1 noticeably decreased after combination treatment, while levels of the pro-apoptotic BH3-only protein BIM increased (Figure 4A-B).

**Figure 4.**
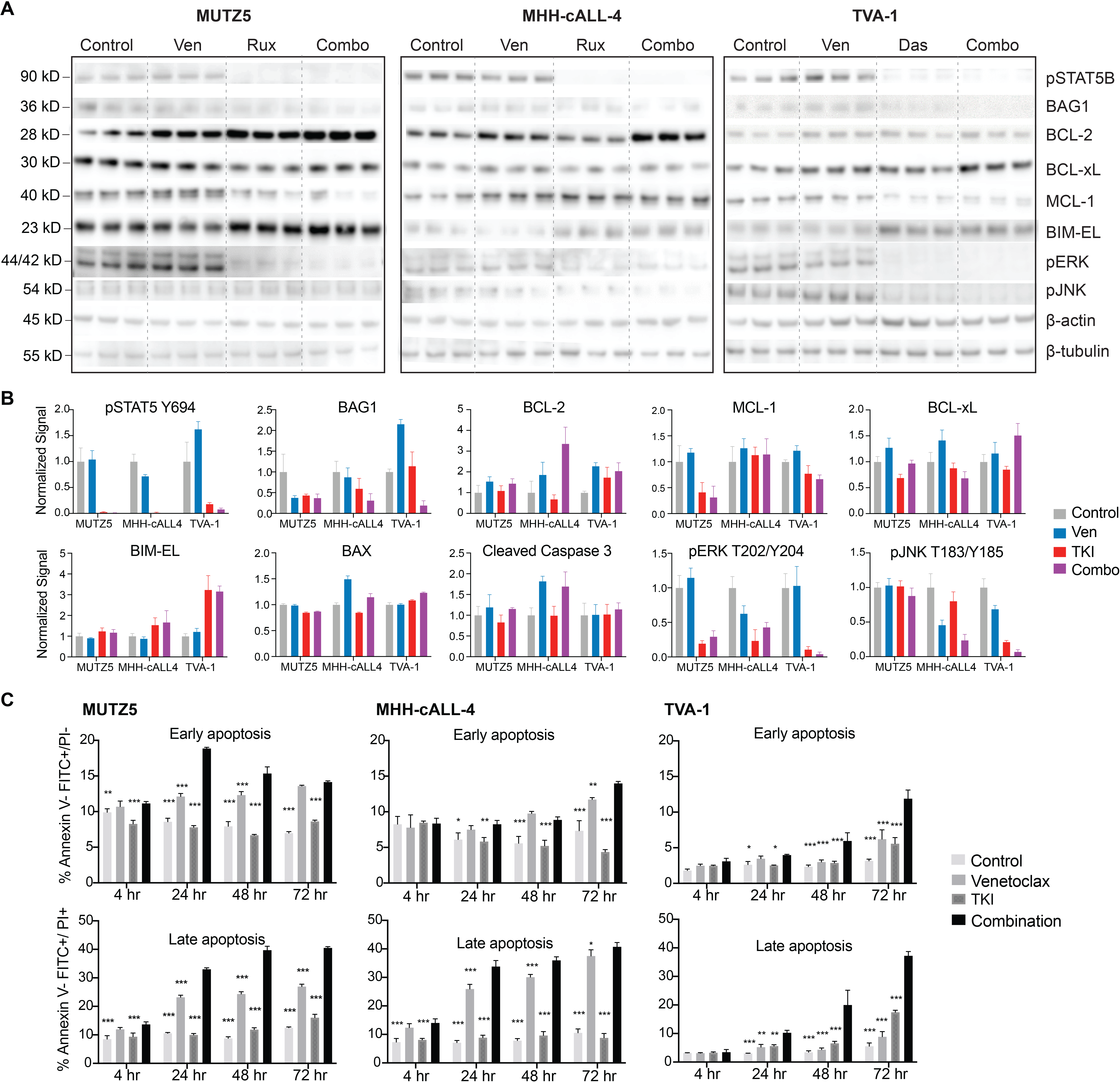
Effects of combined kinase and BCL-2 inhibition on intracellular phosphosignaling, apoptosis proteins, and functional apoptosis. **(A)** Immunoblot images and **(B)** Normalized immunoblotting signal intensities of phosphorylated (p) STAT5, BAG1, BCL-2 family proteins, and pERK and pJNK in Ph-like ALL cell lines treated *in vitro* with venetoclax, TKI (ruxolitinib for MUTZ5 and MHH-cALL-4, dasatinib for TVA-1), or both drugs. Densitometry signals were normalized to β-actin or β-tubulin loading controls and displayed graphically relative to 0.1% DMSO control treatment. Each bar represents mean +/− standard deviation (SD) of 3 technical replicates. BAX and cleaved caspase-3 targets were assessed by apoptosis protein arrays. **(C)** Time course of apoptosis under single or combination drug conditions was assessed by annexin V/propidium iodide (PI) co-staining and flow cytometric analysis. MUTZ5 and MHH-cALL-4 cells were treated with 0.1 μM venetoclax, 1 μM ruxolitinib, both drugs, or 0.1% DMSO control. TVA-1 cells were treated with 50 nM venetoclax, 0.5 nM dasatinib, both drugs, or 0.1% DMSO control. Early apoptosis (assessed by percent annexin V+/PI-, left panels) and late apoptosis/necrosis (assessed by percent annexin V+/PI+, right panels) are shown for each time point. Each bar represents mean ± SD of 3 replicates. Significance of control or single drug as compared to combination indicated by asterisks above bars: *P ≤ 0.05, **P ≤ 0.01, ***P ≤ 0.001 using 2-way ANOVA and Dunnett’s post-test correction.

We next hypothesized that the synergistic decrease in cell viability seen with combined TKI and venetoclax treatment could be due to augmentation of apoptosis. We thus determined the proportion of cells undergoing early apoptosis (Annexin V+/ PI-) and late apoptosis/necrosis (Annexin V+/PI+) and found that combining ruxolitinib or dasatinib with venetoclax led to significantly greater apoptosis than either monotherapy with effects detected within 4-24 hours of drug exposure (Figure 4C). Increased cleaved-caspase 3 was detected in both venetoclax and combination drug-treated cells, consistent with the observed increase in apoptosis at 72 hours in these conditions (Figure 4C, Figure S4). Importantly, despite its strong effects on the expression of apoptosis pathway targets, TKI monotherapy did not appreciably increase apoptosis. Together, these data suggest that the anti-leukemia effects observed from combination drug treatment may be due to TKI-mediated “apoptotic priming” of leukemia cells in decreasing anti-apoptotic proteins while increasing pro-apoptosis proteins, resulting in enhanced apoptosis only when combined with venetoclax.

### Combination of venetoclax and TKI shows potent anti-leukemia efficacy in multiple patient-derived xenograft models of Ph-like ALL and is well-tolerated *in vivo*

To validate pharmacologically our OptiCon-predicted pairing *in vivo*, we investigated the potential efficacy of combined venetoclax and ruxolitinib or dasatinib treatment in six different patient-derived xenograft ( PDX) Ph-like ALL models comprised of *CRLF2*-rearranged or *ABL1*-rearranged genetic backgrounds (Table S7). Given that optimal inhibitor dosing for combination therapy may differ from optimal monotherapy dosing, we tested two dose levels (50% and 100%) of inhibitors in several PDX models to model potential synergy and to assess if anti-leukemia benefit could be achieved with lower dosing.

As hypothesized, we observed significant inhibition of leukemia proliferation in peripheral blood and end-study spleens in most *CRLF2*-rearranged and *ABL1*-rearranged Ph-like ALL PDX models treated with combined venetoclax and ruxolitinib or dasatinib (Figure 5). Detected combination treatment effects were also superior to TKI and/or venetoclax monotherapy in many models. Additionally, near-curative effects of ruxolitinib and venetoclax or dasatinib and venetoclax were observed in end-study spleens of three PDX models (*CRLF2*-rearranged/*JAK2*-mutant JH331, *ABL1*-rearranged NH011 and TVA1). Interestingly, combined TKI and venetoclax at 50% ‘half-dosing’ also had more potent anti-leukemia effects than full monotherapy dosing of either agent, suggesting enhanced STAT5 and BCL-2 co-targeting in these leukemias. While most models did not demonstrate appreciable single-agent venetoclax activity, our surprisingly ruxolitinib-resistant JAK-mutant JH331 PDX model was exquisitely sensitive to BCL-2 inhibition and further showed superior inhibition of leukemia proliferation with dual ruxolitinib/venetoclax treatment, highlighting potential for JAKi resensitization or resistance reversal. Other *CRLF2*-rearranged models (ALL4364, ALL2128) demonstrated marked sensitivity to ruxolitinib with >50% reduction of ALL burden in end-study spleens, although combined treatment with venetoclax did not further augment anti-leukemia response. Importantly, we also observed excellent tolerability of dual inhibitor treatment for up to four weeks in our PDX models with stability of murine weights (Figure S5) and other physical health parameters.

**Figure 5.**
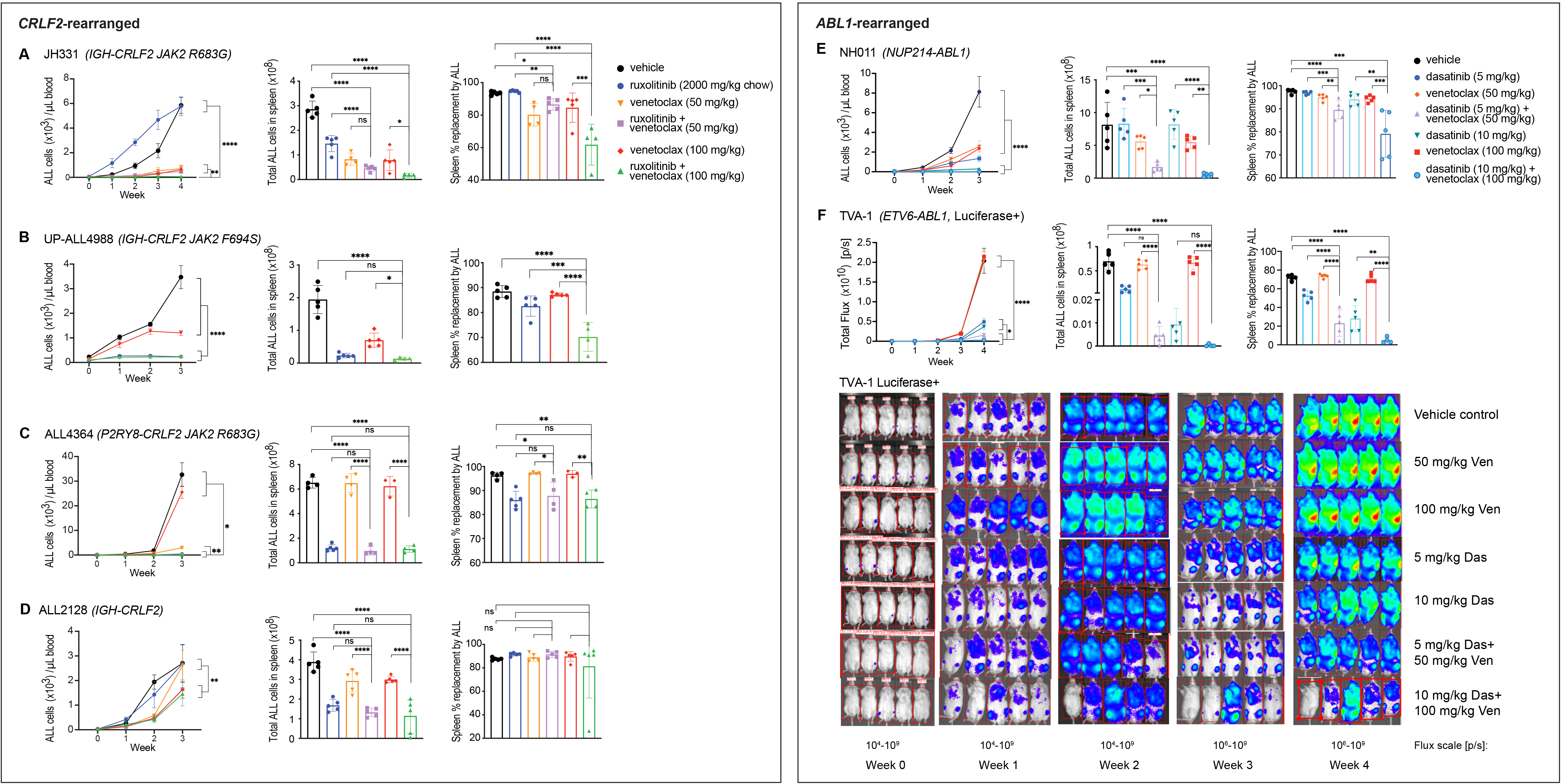
Combined TKI and venetoclax treatment inhibits Ph-like ALL proliferation in vivo. **(A-D)** *CRLF2*-rearranged and **(E-F)** *ABL1*-rearranged Ph-like ALL patient-derived xenograft models (n=5 per treatment arm) were treated with vehicle or inhibitors as delineated above and followed by flow cytometric quantification of human CD10+/CD19+ ALL in murine peripheral blood (left panels) and in end-study spleens (middle and right panels). **(F)** One luciferase-expressing TVA-1 PDX model was followed by bioluminescent imaging (flux measured in photons per second [p/s]) with terminal splenic leukemia burden quantified by flow cytometry as in the other five models. TKI and venetoclax co-treatment significantly inhibited leukemia proliferation in vivo in most PDX models versus inhibitor monotherapy. Error bars represent +/- SD. *P ≤ 0.05, **P ≤ 0.01, ***P ≤ 0.001, ****P ≤ 0.0001 using 2-way ANOVA for peripheral blood analyses over time and 1-way ANOVA for end-study spleen analyses. ns = not significant.

## DISCUSSION

Omics analyses in the study of human cancers often generate large candidate gene lists that can be difficult to prioritize for pharmacologic targeting. We employed an innovative systems biology strategy for unbiased identification of synergistic pathways as candidates for combination therapy. This approach efficiently narrowed the search space of relevant oncogenic pathways and is also broadly applicable. We applied our network-controllability approach to a clinically high-risk leukemia subtype and identified 14 novel optimal control nodes in Ph-like ALL. The largest fraction of the OCNs are involved in kinase signaling, followed by metabolism, transcriptional regulation, splicing, and apoptosis (Figure 1B), suggesting relative dependencies of Ph-like ALL on these biological processes. We focused further experimental validation studies upon the highly ranked OCN pair *STAT5B* and *BAG1* given the strong biologic rationale for this Ph-like ALL-specific OCN pairing. Furthermore, there is pragmatic translational potential for investigating combination therapy using the TKIs ruxolitinib or dasatinib and the BCL-2 inhibitor venetoclax given their clinical availability and established adult and pediatric dosing. Venetoclax has not been extensively investigated in Ph-like ALL, although several preclinical studies have reported preliminary efficacy of combining BCL-2 inhibitors with TKIs in other ALL types such *BCR-ABL1*+ models^47^ and *IL7R*-mutant T-ALL models^48^. OptiCon’s nomination of *STAT5B* and *BCL2*-related pathways from integrative-omics analysis of primary Ph-like ALL samples provides independent rationale for co-targeting these specifically in the Ph-like ALL subtype.

Intriguingly, we demonstrated that ‘precision drugging’ of the *STAT5B* and *BAG1* pair of synergistic regulators altered the transcriptomic state of Ph-like ALL to become more similar to other chemosensitive B-ALL subtypes. These results support the basis of network controllability theory that successful identification of key control nodes can shift cells from one transcriptomic state to another desired one via specific perturbation. Although the transcriptomic effects of joint BCL-2 and kinase inhibition appear to be driven in large part by the TKI, the phenotypic effects of combined therapy in enhancing cell death seem driven by venetoclax. Our functional assays of apoptosis and cell viability combined with transcriptional and protein analyses suggest that TKI monotherapy may be priming Ph-like cells for apoptosis by altering the fine balance between pro-apoptotic and anti-apoptotic protein expression.

OptiCon analysis also elucidated potential crosstalk pathways that mediate synergy between the STAT5 signaling and BCL-2 pathways in Ph-like ALL. We found that several MAPK pathway genes (*e.g.*, *MAPK8* (JNK1) and *MAPK11* (p38)) participate in crosstalk between the OptiCon-predicted OCNs *STAT5B* and *BAG1*. While mutations in MAPK pathway genes have been reported in Ph-like ALL^7^, the mechanistic relevance of deregulated MAPK signaling in Ph-like ALL is not well-understood. JNKs are known to phosphorylate and regulate BCL-2, BIM, and BAD with effects on apoptosis in a context and cell-type dependent manner, and ERK1/2 activation has been associated with anti-apoptotic effects^49,50^. Our identification of their negative regulator *DUSP3* as an OCN and our surprising findings that co-treatment with venetoclax and ruxolitinib or dasatinib also decreased phosphorylated ERK1/2 and JNK levels suggest that MAP kinases are involved in critical Ph-like ALL signaling crosstalk.

Our studies reveal in an unbiased manner that both *CRLF2*-rearranged and *ABL1*-rearranged Ph-like ALL appear to have previously-unknown BCL-2 dependencies. Our additional demonstration of superior *in vivo* anti-leukemia effects of combining ruxolitinib or dasatinib and venetoclax in multiple PDX models with various genetic backgrounds, provides strong preclinical rationale for bench-to-bedside development of dual BCL-2 and kinase inhibition in next-generation clinical trials for patients with Ph-like ALL. Further elucidation of key genetic and molecular factors that may contribute to the observed heterogeneity of inhibitor treatment responses in our Ph-like ALL PDX models will facilitate potential future clinical translation of these findings. Importantly, we also observed effective *in vivo* leukemia burden reduction when venetoclax and dasatinib were used at subtherapeutic dosing, suggesting the ability to achieve effective anti-leukemia activity while potentially reducing therapy-associated toxicity. This observation lends additional support for our network-based approach which takes pathway crosstalk into account to spare unperturbed pathways in order to minimize potential toxicity.

Collectively, our application of a systems biology framework to a high-risk leukemia subtype provides critical new insights regarding cancer gene network controllability and the ability to facilitate unbiased discovery of novel target pairings. While our results certainly provide compelling rationale for clinical investigation of new dual venetoclax and ruxolitinib or dasatinib strategies for patients with Ph-like ALL, our network controllability-based methodology for inferring synergistic gene regulatory nodes also provides an innovative paradigm for rational design of combination therapy approaches that has wide applicability to other human cancers and diseases.

## ACKNOWLEDGEMENTS

This work was supported by the National Institutes of Health (NIH)/National Child Health and Human Development award T32HD043021 to the Children’s Hospital of Philadelphia (YYD), NIH/National Cancer Institute awards K12CA076931 to the University of Pennsylvania (YYD), K08CA184418 (SKT), U01CA232486 (SKT), and U01CA243072 (SKT, KT), Department of Defense Translational Team Science award CA180683P1 (SKT), the V Foundation for Cancer Research (SKT), and institutional support from the Children’s Hospital of Philadelphia Center for Childhood Cancer Research (CHOP CCCR) (SKT, KT) and CHOP Research Institute (YYD). We acknowledge the Children’s Oncology Group and the CHOP CCCR biorepositories for provision of primary patient leukemia specimens. We also thank Dr David Fruman at the University of California, Irvine for kindly sharing the TVA-1 patient-derived xenograft model and Dr Matthew Stubbs at the Incyte Corporation for generous provision of ruxolitinib rodent chow for animal studies.

## AUTHOR CONTRIBUTIONS

YD, SKT, and KT conceived the study. YD designed and performed experiments, analyzed and interpreted data, and wrote the manuscript. HK, GMC, and YH developed analytic pipelines and performed data analysis. KM, JPL, and DHA performed experiments and analyzed data. RZ, JX, and YH performed data analysis. SKT and KT supervised the overall study, designed experiments, analyzed and interpreted data, and wrote the manuscript. All authors approved the final version of the manuscript.

## REFERENCES

1. Garraway LA, Jänne PA. Circumventing cancer drug resistance in the era of personalized medicine. Cancer Discov. 2012;2(3):214–226.

2. Huang A, Garraway LA, Ashworth A, Weber B. Synthetic lethality as an engine for cancer drug target discovery. Nat. Rev. Drug Discov. 2020;19(1):23–38.

3. Laufer C, Fischer B, Billmann M, Huber W, Boutros M. Mapping genetic interactions in human cancer cells with RNAi and multiparametric phenotyping. Nature Methods. 2013;10(5):427–431.

4. Du D, Roguev A, Gordon DE, et al. Genetic interaction mapping in mammalian cells using CRISPR interference. Nat. Methods. 2017;14(6):577–580.

5. Den Boer ML, van Slegtenhorst M, De Menezes RX, et al. A subtype of childhood acute lymphoblastic leukaemia with poor treatment outcome: a genome-wide classification study. Lancet Oncol. 2009;10(2):125–134.

6. Mullighan CG, Zhang J, Harvey RC, et al. JAK mutations in high-risk childhood acute lymphoblastic leukemia. Proc. Natl. Acad. Sci. U. S. A. 2009;106(23):9414–9418.

7. Roberts KG, Li Y, Payne-Turner D, et al. Targetable Kinase-Activating Lesions in Ph-like Acute Lymphoblastic Leukemia. N. Engl. J. Med. 2014;371(11):1005–1015.

8. Reshmi SC, Harvey RC, Roberts KG, et al. Targetable kinase gene fusions in high-risk B-ALL: a study from the Children’s Oncology Group. Blood. 2017;129(25):3352–3361.

9. Jain N, Roberts KG, Jabbour E, et al. Ph-like acute lymphoblastic leukemia: a high-risk subtype in adults. Blood. 2017;129(5):572–581.

10. Tasian SK, Loh ML, Hunger SP. Philadelphia chromosome-like acute lymphoblastic leukemia. Blood. 2017;130(19):2064–2072.

11. Harvey RC, Mullighan CG, Wang X, et al. Identification of novel cluster groups in pediatric high-risk B-precursor acute lymphoblastic leukemia with gene expression profiling: correlation with genome-wide DNA copy number alterations, clinical characteristics, and outcome. Blood. 2010;116(23):4874–4884.

12. Maude SL, Tasian SK, Vincent T, et al. Targeting JAK1/2 and mTOR in murine xenograft models of Ph-like acute lymphoblastic leukemia. Blood. 2012;120(17):3510–3518.

13. Tasian SK, Teachey DT, Li Y, et al. Potent efficacy of combined PI3K/mTOR and JAK or ABL inhibition in murine xenograft models of Ph-like acute lymphoblastic leukemia. Blood. 2016;129(2):177–187.

14. Ding YY, Stern JW, Jubelirer TF, et al. Clinical efficacy of ruxolitinib and chemotherapy in a child with Philadelphia chromosome-like acute lymphoblastic leukemia with GOLGA5-JAK2 fusion and induction failure. Haematologica. 2018;103(9):e427–e431.

15. Tasian SK, Doral MY, Borowitz MJ, et al. Aberrant STAT5 and PI3K/mTOR pathway signaling occurs in human CRLF2-rearranged B-precursor acute lymphoblastic leukemia. Blood. 2012;120(4):833–842.

16. Hu Y, Chen C-H, Ding Y-Y, et al. Optimal control nodes in disease-perturbed networks as targets for combination therapy. Nat. Commun. 2019;10(1):2180.

17. Liu Y-Y, Slotine J-J, Barabási A-L. Controllability of complex networks. Nature. 2011;473(7346):167–173.

18. Ma X, Liu Y, Liu Y, et al. Pan-cancer genome and transcriptome analyses of 1,699 paediatric leukaemias and solid tumours. Nature. 2018;555(7696):371.

19. Gotesman M, Vo T-TT, Herzog L-O, et al. mTOR inhibition enhances efficacy of dasatinib in ABL-rearranged Ph-like B-ALL. Oncotarget. 2018;9(5):6562–6571.

20. Chou T-C. Drug Combination Studies and Their Synergy Quantification Using the Chou-Talalay Method. Cancer Res. 2010;70(2):440–446.

21. Bhojwani D, Pei D, Sandlund JT, et al. ETV6-RUNX1-positive childhood acute lymphoblastic leukemia: improved outcome with contemporary therapy. Leukemia. 2012;26(2):265–270.

22. Heerema NA, Sather HN, Sensel MG, et al. Prognostic Impact of Trisomies of Chromosomes 10, 17, and 5 Among Children With Acute Lymphoblastic Leukemia and High Hyperdiploidy (> 50 Chromosomes). J. Clin. Orthod. 2000;18(9):1876–1887.

23. Zhang J, McCastlain K, Yoshihara H, et al. Deregulation of DUX4 and ERG in acute lymphoblastic leukemia. Nat. Genet. 2016;48(12):1481–1489.

24. Matsumoto A, Masuhara M, Mitsui K, et al. CIS, a cytokine inducible SH2 protein, is a target of the JAK-STAT5 pathway and modulates STAT5 activation. Blood. 1997;89(9):3148–3154.

25. Hurtz C, Wertheim GB, Loftus JP, et al. Oncogene-independent BCR-like signaling adaptation confers drug resistance in Ph-like ALL. J. Clin. Invest. 2020;

26. Aveic S, Viola G, Accordi B, et al. Targeting BAG-1: A novel strategy to increase drug efficacy in acute myeloid leukemia. Exp. Hematol. 2015;43(3):180–190.e6.

27. Amand M, Erpicum C, Bajou K, et al. DUSP3/VHR is a pro-angiogenic atypical dual-specificity phosphatase. Mol. Cancer. 2014;13:108.

28. Bride KL, Vincent TL, Im S-Y, et al. Preclinical efficacy of daratumumab in T-cell acute lymphoblastic leukemia. Blood. 2018;131(9):995–999.

29. Jee HJ, Kim AJ, Song N, et al. Nek6 overexpression antagonizes p53-induced senescence in human cancer cells. Cell Cycle. 2010;9(23):4703–4710.

30. Zhu F, Shi Z, Qin C, et al. Therapeutic target database update 2012: a resource for facilitating target-oriented drug discovery. Nucleic Acids Res. 2012;40(Database issue):D1128–36.

31. Wishart DS, Feunang YD, Guo AC, et al. DrugBank 5.0: a major update to the DrugBank database for 2018. Nucleic Acids Res. 2018;46(D1):D1074–D1082.

32. Cotto KC, Wagner AH, Feng Y-Y, et al. DGIdb 3.0: a redesign and expansion of the drug--gene interaction database. Nucleic Acids Res. 2018;46(D1):D1068–D1073.

33. Donato MD, Righino B, Filippetti F, et al. Identification and antitumor activity of a novel inhibitor of the NIMA-related kinase NEK6. Sci. Rep. 2018;8(1):16047.

34. Tasian SK, Assad A, Hunter DS, Du Y, Loh ML. A Phase 2 Study of Ruxolitinib with Chemotherapy in Children with Philadelphia Chromosome-like Acute Lymphoblastic Leukemia (INCB18424-269/AALL1521): Dose-Finding Results from the Part 1 Safety Phase. Blood. 2018;132(Supplement 1):555–555.

35. Aveic S, Pigazzi M, Basso G. BAG1: The Guardian of Anti-Apoptotic Proteins in Acute Myeloid Leukemia. PLoS One. 2011;6(10):e26097.

36. Takayama S, Bimston DN, Matsuzawa S, et al. BAG-1 modulates the chaperone activity of Hsp70/Hsc70. EMBO J. 1997;16(16):4887–4896.

37. Stilgenbauer S, Eichhorst B, Schetelig J, et al. Venetoclax in relapsed or refractory chronic lymphocytic leukaemia with 17p deletion: a multicentre, open-label, phase 2 study. The Lancet Oncology. 2016;17(6):768–778.

38. Pollyea DA, Amaya M, Strati P, Konopleva MY. Venetoclax for AML: changing the treatment paradigm. Blood Adv. 2019;3(24):4326–4335. Blood Adv. 2020;4(6):1020.

39. Weigert O, Lane AA, Bird L, et al. Genetic resistance to JAK2 enzymatic inhibitors is overcome by HSP90 inhibition. J. Exp. Med. 2012;209(2):259–273.

40. Zhou T, Georgeon S, Moser R, et al. Specificity and mechanism-of-action of the JAK2 tyrosine kinase inhibitors ruxolitinib and SAR302503 (TG101348). Leukemia. 2014;28(2):404–407.

41. McInnes L, Healy J. UMAP: Uniform Manifold Approximation and Projection for Dimension Reduction. 2018;

42. Winters AC, Bernt KM. MLL-Rearranged Leukemias—An Update on Science and Clinical Approaches. Frontiers in Pediatrics. 2017;5:4.

43. Kanwal Z, Zakrzewska A, den Hertog J, et al. Deficiency in hematopoietic phosphatase ptpn6/Shp1 hyperactivates the innate immune system and impairs control of bacterial infections in zebrafish embryos. J. Immunol. 2013;190(4):1631–1645.

44. Andreeff M, Jiang S, Zhang X, et al. Expression of Bcl-2-related genes in normal and AML progenitors: changes induced by chemotherapy and retinoic acid. Leukemia. 1999;13(11):1881–1892.

45. Tessoulin B, Papin A, Gomez-Bougie P, et al. BCL2-Family Dysregulation in B-Cell Malignancies: From Gene Expression Regulation to a Targeted Therapy Biomarker. Front. Oncol. 2019;8:645.

46. Yang-Yen H-F. Mcl-1: a highly regulated cell death and survival controller. J. Biomed. Sci. 2006;13(2):201–204.

47. Leonard JT, Rowley JSJ, Eide CA, et al. Targeting BCL-2 and ABL/LYN in Philadelphia chromosome–positive acute lymphoblastic leukemia. Sci. Transl. Med. 2016;8(354):354ra114–354ra114.

48. Senkevitch E, Li W, Hixon JA, et al. Inhibiting Janus Kinase 1 and BCL-2 to treat T cell acute lymphoblastic leukemia with IL7-Rα mutations. Oncotarget. 2018;9(32):22605–22617.

49. Yue J, López JM. Understanding MAPK Signaling Pathways in Apoptosis. Int. J. Mol. Sci. 2020;21(7):.

50. Dhanasekaran DN, Reddy EP. JNK-signaling: A multiplexing hub in programmed cell death. Genes Cancer. 2017;8(9–10):682–694.

51. He B, Gao P, Ding Y-Y, et al. Diverse noncoding mutations contribute to deregulation of cis-regulatory landscape in pediatric cancers. Sci Adv. 2020;6(30):eaba3064.

52. Kanehisa M, Furumichi M, Tanabe M, Sato Y, Morishima K. KEGG: new perspectives on genomes, pathways, diseases and drugs. Nucleic Acids Res. 2017;45(D1):D353–D361.

53. Fabregat A, Sidiropoulos K, Garapati P, et al. The Reactome pathway Knowledgebase. Nucleic Acids Res. 2016;44(D1):D481–7.

54. Schaefer CF, Anthony K, Krupa S, et al. PID: the Pathway Interaction Database. Nucleic Acids Res. 2009;37(Database issue):D674–9.

55. Ritchie ME, Phipson B, Wu D, et al. limma powers differential expression analyses for RNA-sequencing and microarray studies. Nucleic Acids Res. 2015;43(7):e47.

56. Dobin A, Davis CA, Schlesinger F, et al. STAR: ultrafast universal RNA-seq aligner. Bioinformatics. 2013;29(1):15–21.

57. Trapnell C, Williams BA, Pertea G, et al. Transcript assembly and quantification by RNA-Seq reveals unannotated transcripts and isoform switching during cell differentiation. Nat. Biotechnol. 2010;28(5):511–515.

58. Liao Y, Smyth GK, Shi W. featureCounts: an efficient general purpose program for assigning sequence reads to genomic features. Bioinformatics. 2014;30(7):923–930.

59. Law CW, Chen Y, Shi W, Smyth GK. Voom: precision weights unlock linear model analysis tools for RNA-seq read counts. Genome Biol. 2014;15(2):R29.

60. Huang DW, Sherman BT, Lempicki RA. Systematic and integrative analysis of large gene lists using DAVID bioinformatics resources. Nat. Protoc. 2009;4(1):44–57.

61. Johnson WE, Li C, Rabinovic A. Adjusting batch effects in microarray expression data using empirical Bayes methods. Biostatistics. 2007;8(1):118–127.

62. Barbie DA, Tamayo P, Boehm JS, et al. Systematic RNA interference reveals that oncogenic KRAS-driven cancers require TBK1. Nature. 2009;462(7269):108–112.

63. Fischer U, Forster M, Rinaldi A, et al. Genomics and drug profiling of fatal TCF3-HLF−positive acute lymphoblastic leukemia identifies recurrent mutation patterns and therapeutic options. Nat. Genet. 2015;47(9):ng.3362.

64. Gill S, Tasian SK, Ruella M, et al. Preclinical targeting of human acute myeloid leukemia and myeloablation using chimeric antigen receptor-modified T cells. Blood. 2014;123(15):2343–2354. Blood. 2016;128(21):2585.

